# Can functional traits explain phylogenetic signal in the composition of a plant community?

**DOI:** 10.1101/032938

**Authors:** Daijiang Li, Anthoy R. Ives, Donald M. Waller

**Author notes:** Correspondence: Daijiang Li. **Authorship:** DL and AI designed the study and performed the analyses. DL collected the vegetation and environmental data. DL and DW collected the functional trait data. DL wrote the first draft of the manuscript, and all authors worked together on revisions.

## Abstract

Phylogeny-based and functional trait-based analyses are used widely to study community composition. In principle, knowing all information about species traits should completely explain phylogenetic patterns in community composition. In reality, phylogenies may contain more information than the collection of measured traits. The extent to which functional trait information makes phylogenetic information redundant, however, is unknown. We used phylogenetic linear mixed models to analyze community composition of 55 understory plant species distributed across 30 forest sites in central Wisconsin. These communities showed strong phylogenetic attraction. Most of the 15 measured functional traits showed strong phylogenetic signal, but they only reduced the strength of phylogenetic community patterns in the abundances and presence/absences of co-occurring species by 57% and 89%, respectively, falling short of fully explaining phylogenetic community structure. Our study demonstrates the value of phylogenies in studying of community composition, especially with abundance data, even when rich functional trait data are available.

## Introduction

Functional traits, arising as innovations through evolution, can capture essential aspects of species’ morphology, ecophysiology, and life-history strategy (McGill *et al*. 2006; Violle *et al*. 2007). Although closely related species can differ greatly in some functional traits due to rapid evolution or ecological convergence (Losos, 2008, 2011), most functional traits show strong phylogenetic signal (Freckleton *et al*. 2002; Webb *et al*. 2002, Moles *et al*. 2005, Donoghue 2008). Functional traits, with or without phylogenetic signal, are known to influence the species composition of communities, thereby providing mechanistic links between fundamental ecological processes and community structure (McGill *et al*. 2006; Violle *et al*. 2007; Adler *et al*. 2013). Functional traits also provide a common currency that facilitates comparisons among species and across regions, allowing us to assess the generality of patterns and predictions in community ecology (McGill *et al*. 2006). This has lead to a proliferation of studies using functional traits to understand community composition. Functional trait-based approaches, however, are limited by the fact that it is impossible to measure all potentially important functional traits affecting the distribution of species.

Even in the absence of functional trait information, it is still possible to infer the effects of (unmeasured) functional traits on community composition by investigating phylogenetic patterns in community composition. Phylogenies play an important role in community ecology by giving information about evolutionary relationships among species (Graves & Gotelli, 1993; Losos 1996; Baum & Smith, 2012). Because phylogenetically related species often share similar functional trait values, we expect phylogenetically related species to co-occur more often in the same communities reflecting their shared environmental tolerances. Conversely, if phylogenetically related species have similar traits that cause them to compete with each other, then closely related species may be less likely to co-occur. These and other processes relating functional traits to community composition likely lead to phylogenetic signatures in how species are distributed among communities (Webb *et al*. 2002). However, in principle, if we have information for all relevant functional traits, then we expect phylogeny to provide little additional information relevant for community composition. That is, when all of the functional traits affecting community composition are known, we do not expect the unexplained residual variation in the occurrence of species to have phylogenetic signal (Ives & Helmus, 2011).

In practice, we cannot obtain information about all relevant functional traits. In addition, phylogenetic signals in community composition may result from factors beyond functional traits, such as the biogeographical patterns generated as species disperse across a landscape (Ricklefs *et al*. 1993; Moen *et al*. 2009). If these forces are important, then even after accounting for all functional traits whose measurements are available, we should expect phylogenies to contain additional information about community composition (Vane-Wright *et al*. 1991; Cadotte *et al*. 2009). Thus far, however, we are aware of no study that has explicitly assessed the overlap between information from traits versus phylogeny. Here, we ask how much of the phylogenetic signal in the composition of a plant community assemblage can be explained by functional traits (Fig. 1).

**Figure 1.**
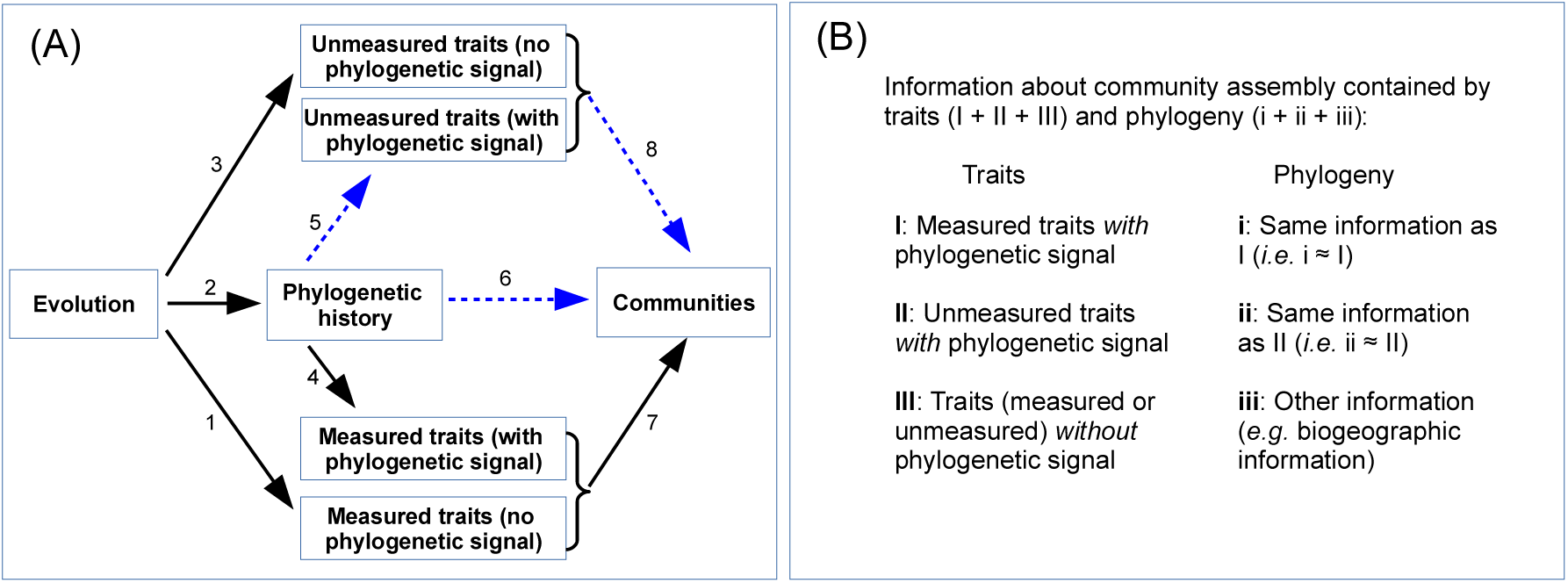
Schematic diagram of the conceptual framework of the study. (A) Evolution is the ultimate source of all trait values, although only some traits have phylogenetic signal that reflects phylogenetic history (arrows 2, 4 and 5). Other traits do not (arrows 1 and 3), possibly because these traits evolve rapidly or experience convergent evolution. Community composition is determined by unmeasured and measured traits, and also by additional processes that could generate phylogenetic signal, such as biogeographical patterns in the distribution of species. Phylogenetic patterns in community composition can be generated from measured and unmeasured traits with phylogenetic signal (arrows 7 and 8), and by other phylogenetic processes (arrow 6). The question we address is how much of the phylogenetic signal in community composition can be explained by measured functional traits, and whether after accounting for these traits there is residual phylogenetic signal that could have been generated by unmeasured traits or other phylogenetic processes. (B) Traits and phylogeny contain overlapping and complementary information about how communities are assembled. Here, we focus on estimating the proportion of this overlapping information that the phylogeny contains (i.e., the magnitude of *i* relative to *i* + *ii* + *iii*). Note that we do not try to explain the proportion of overlapping information that functional traits contain (i.e., the magnitude of *I* relative to *I* + *II*+ *III)* due to our inability to estimate the amount of information provided by unmeasured traits and hence estimate (*I* + *II* + *III)*.

We analyzed data on the abundance of 55 understory plant species distributed across 30 Wisconsin pine barrens sites (Li & Waller 2015). For each species, we had data on 15 functional traits and a recent highly resolved phylogeny (Cameron *et al. unpublished manuscript*^1^). At each site, we measured 20 environmental variables. Below, we first investigate whether there is phylogenetic pattern in community composition, using a phylogenetic community mixed model that tests for both “phylogenetic attraction” (phylogenetically related species more likely to occur in the same communities) and “phylogenetic repulsion.” If there is phylogenetic pattern, then it could be produced by measured functional traits that themselves have phylogenetic signal (Fig. 1, arrows 2, 4, and 7), unmeasured functional traits with phylogenetic signal (Fig. 1, arrows 2, 5, and 8), or phylogenetic processes unrelated to functional traits (Fig. 1, arrow 6). We then developed a phylogenetic community mixed model incorporating the measured functional traits to ask whether there is phylogenetic signal in the residual variation in community composition after the effects of these traits are removed. This analysis tests the hypothesis that we can explain all of the phylogenetic pattern in community composition using measured functional traits. Finally, we use a phylogenetic community mixed model to investigate whether phylogenetically related species respond similarly to environmental gradients across the communities. The motivation for this final analysis is to indirectly identify possible unmeasured functional traits that might play a role in community assembly. In cases where phylogenetically related species respond similarly to an environmental gradient, species presumably share traits that confer similar tolerances to, or preferences for, specific environmental conditions. Thus, this final analysis could point towards additional functional traits that might be relevant for explaining patterns in community composition.

**Figure 2:**
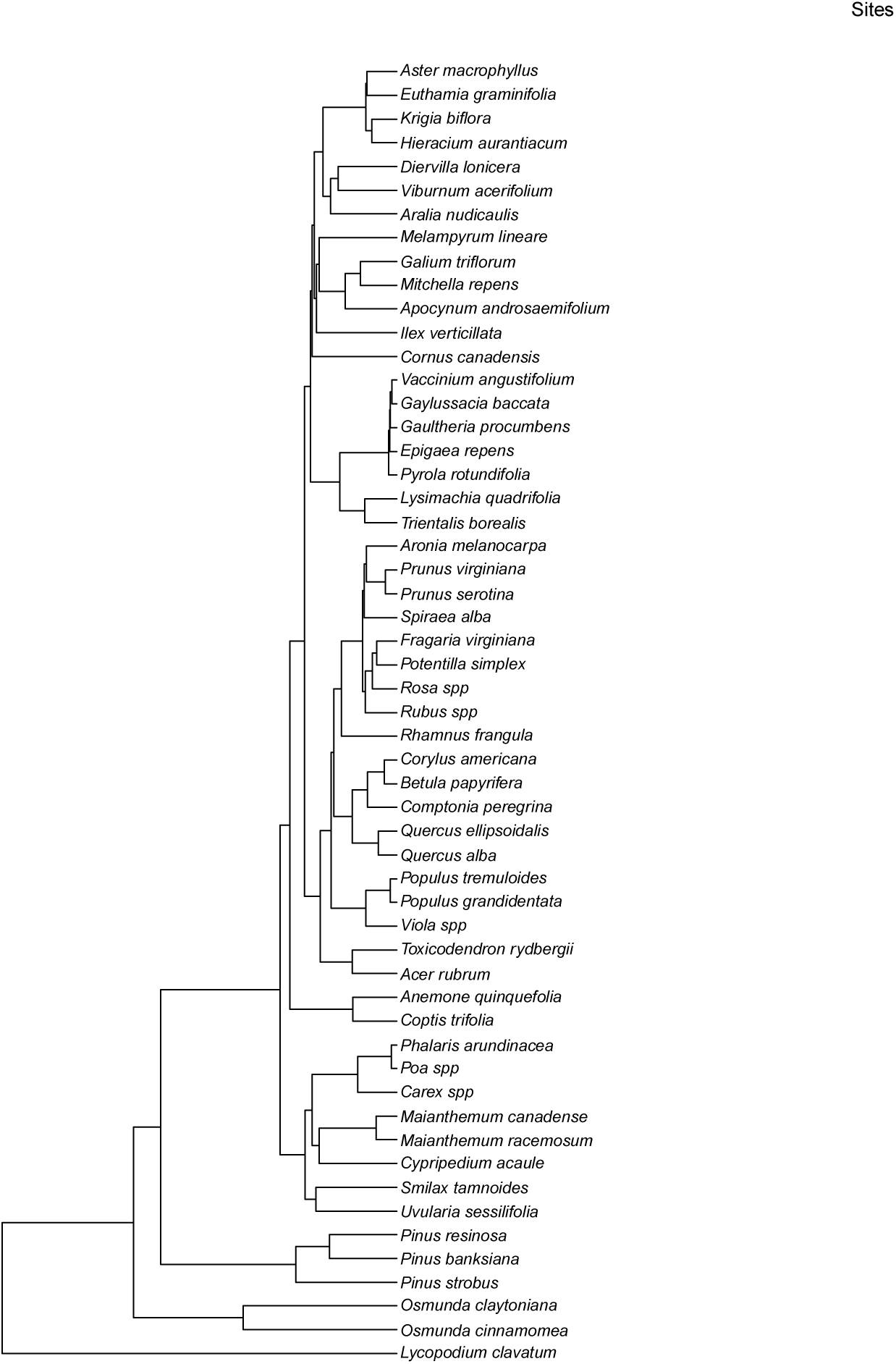
Phylogeny and relative abundance of the 55 common plant species found in the pine barrens of central Wisconsin in 2012. The area of dots is proportional to abundances within each site.

## Methods

### Data

*Community composition*. – We sampled 30 pine barrens forest sites in the central Wisconsin sand plains in 2012 using 50 1-*m*^2^ quadrats placed along five transects at each site. Within each quadrat, we recorded the presence/absence of all understory vascular plant species (see Li & Waller 2015 for details). Across all sites, we recorded 152 species. For the analyses other than the initial exploration of phylogenetic patterns in community composition, we focused on the 55 species that occurred in three or more communities. We did this because we did not have functional trait data for many rare species, and we also wanted to limit the number of zeros in the data set.

*Functional traits*. – For the 55 focal species, we measured 11 continuous and four categorical functional traits on at least 12 individuals (four from each of at least three populations) using standard protocols (Pérez-Harguindeguy *et al*. 2013). Continuous traits include seed mass (g/seed), plant height *(cm)*, specific leaf area (SLA, *m^2^/kg)*, leaf dry matter content (LDMC, %), leaf circularity (dimensionless), leaf length (*cm*), leaf width (cm), leaf thickness (*mm*), leaf carbon concentration (%), leaf nitrogen concentration (%), and stem dry matter content (SDMC, %). We aggregated categories of each categorical trait into two levels: growth form (woody vs. non-woody), life cycle (annual vs. non-annual), and pollination mode (biotic vs. abiotic). We divided seed dispersal mode into three binary variables (wind dispersed vs. not, animal dispersed vs. not, and unassisted vs. assisted dispersal). Collectively, these functional traits, covering the leaf-height-seed (LHS) plant ecology strategy (Westoby, 1998), represent multidimensional functions of plants associated with resource use, competitive ability, dispersal ability, etc. For analyses, we log-transformed highly skewed traits first and then Z-transformed the trait values to have means of zero and standard deviations of one, allowing coefficients in the mixed models to be interpreted as effect sizes.

*Phylogeny*. – The phylogeny used in this study is a subset of a phylogeny for all vascular plants in Wisconsin (Cameron *et al. unpublished manuscript)*. Briefly, Cameron *et al*. used two plastid DNA barcode loci *rbcL* and *matK* to build the phylogeny using maximum likelihood (ML) in the program R_AXML_ (Stamatakis, 2014). The phylogeny was then time-calibrated using the branch length adjuster *(bladj)* available in the program phylocom (Webb *et al*. 2008).

*Environmental data*. – At each site, we pooled six soil samples to measure the soil properties listed in Table 4. We also took six vertical fish-eye photographic images at each site to measure canopy cover. To characterize climatic conditions, we extracted daily precipitation and minimum temperature for each site from interpolated values estimated by Kucharik *et al*. (2010) from 2002 to 2006 (data after 2006 were not available). All environmental variables were Z-transformed.

### Phylogenetic community composition

We performed all analyses using both species abundances and species presence/absences among communities. In the main text we present the analyses of abundance data, because including abundance data in phylogenetic community analyses provides more information about community assembly (Freilich & Connolly, 2015). In the Appendix we present the results for presence/absence data.

We first tested for phylogenetic community structure without including environmental or functional trait information. We used traditional metrics and randomization tests (i.e., null models) to identify whether there was phylogenetic pattern (phylogenetic attraction or repulsion) in the composition of our 30 communities. Specifically, we measured the phylogenetic structure of species abundances at each site using phylogenetic species evenness (PSE, Helmus *et al*. 2007) and mean phylogenetic distance (MPD, Webb, 2000). For each site, we calculated PSE and MPD, and then calculated the mean of these metrics (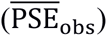 and 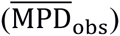) across all 30 sites. To test for phylogenetic pattern, we permuted species randomly among sites (SIM2 in Gotelli, 2000) 4999 times and then calculated metrics base on each permutation data set. If 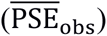 or 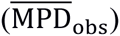 falls below (or above) 97.5% of the permutation values, then we infer a statistically significant phylogenetic attraction (or repulsion). This null permutation model retains the prevalence of each species across sites, but allows sites to change in species richness. Using this null model where sites can vary in species richness is justified, because under the null hypothesis of no phylogenetic signal, the values of PSE and MPD are independent of species richness at the sites. We also performed permutation tests on the presence/absence of species from the 30 sites using phylogenetic species variation (PSV, Helmus *et al*. 2007) and MPD.

In addition to these permutation tests, we fit a phylogenetic linear mixed model (PLMM) to test for phylogenetic community patterns in species abundances. A PLMM establishes a flexible statistical base to subsequently incorporate functional trait and environmental variables. Furthermore, PLMMs tend to have greater statistical power than permutation tests (Ives & Helmus, 2011). To build the PLMM, let *n* be the number of species distributed among *m* sites. Letting *Y* be the *mn* × 1 vector containing the abundance of species *j (j* = 1, …, *n*) at site *s (s* = 1, …, *m*), the PLMM is

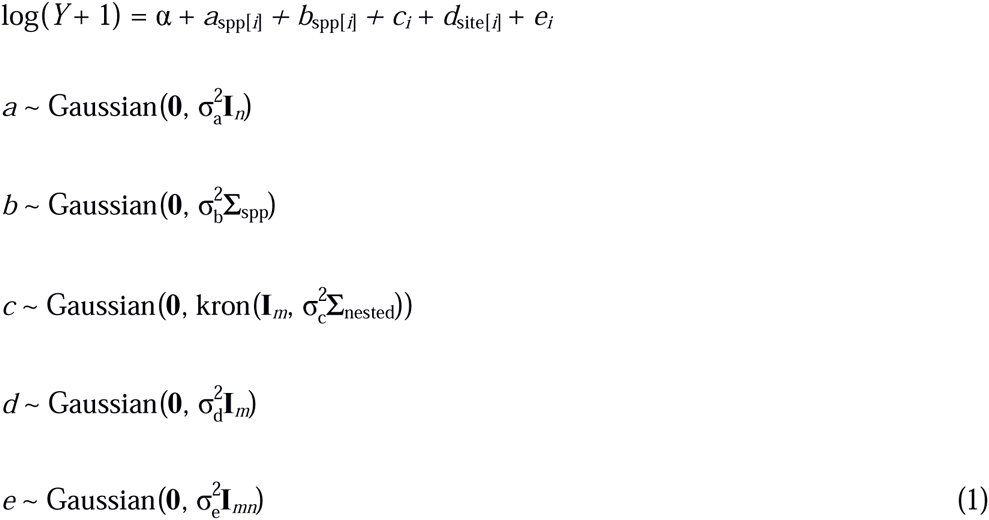

We use the convention of multilevel models here (Gelman & Hill, 2007), with fixed and random effects given by Greek and Latin letters, respectively. The function spp[*i*] maps the observation *i* in vector *Y* to the identity of the species (Gelman & Hill, 2007, p251–252), so *i* takes values from 1 to *mn*. The intercept *α* estimates the overall average log abundance of species across all sites. The following three random variables *a*_spp[_*_i_*_]_, *b*_spp[_*_i_*_]_ and *c_i_* incorporate variation in abundance among plant species. Specifically, the *n* values of *a*_spp[_*_i_*_]_ give differences among species in mean log abundance across all sites and are assumed to be drawn independently from a Gaussian distribution with mean 0 and variance 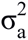 The *n* values of *b*_spp[_*_i_*_]_ also give differences in mean log abundance across sites but are assumed to be drawn from a multivariate Gaussian distribution with covariance matrix 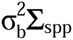, where the *n* × *n* matrix Σ_spp_ is derived from the phylogeny (see next paragraph), and the scalar 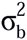 dictates the overall strength of the phylogenetic signal. Thus, *a*_spp[_*_i_*_]_ and *b*_spp[_*_i_*_]_ together capture variation in mean species log abundances that is either unrelated to phylogeny or has phylogenetic signal. The random variable *c_i_* accounts for covariance in the log abundances of plant species nested within sites (using the Kronecker product, kron). Specifically, *c_i_* assesses whether phylogenetically related plant species are more or less likely to co-occur at the same sites. Hence, 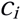 is used to measure either phylogenetic attraction or phylogenetic repulsion; because 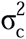 dictates the overall strength of these phylogenetic patterns, it is the key term we are interested in. Random effect *d*_site[_*_i_*_]_ is assumed to contain *m* values, one for each site, that are distributed by a Gaussian distribution with variance 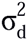 to account for differences in the average log abundances of species from site to site. Finally, *e_i_* captures residual variance 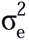.

We derived the phylogenetic covariance matrix Σ_spp_ from the assumption of Brownian motion evolution. If a continuous-valued trait evolves up a phylogenetic tree with a constant probability of slight increases or decreases, the covariance in trait values between two species will be proportional to the length of shared evolution, given by the distance on the phylogenetic tree between the root and the species’ most recent common ancestor (Martins & Hanson 1997). This gives a direct way to convert the phylogeny into a hypothesis about the covariance matrix. For the assessment of phylogenetic attraction within sites, *c_i_*, we use Σ_nested_ = Σ_spp_. For phylogenetic repulsion, we use the matrix inverse of Σ_spp_, Σ_nested_ = (Σ_spp_)^−1^. Theoretical justification for Σ_nested_ = (Σ_spp_)^−1^ comes from a model of competition among community members (Ives & Helmus 2011, Appendix A). Briefly, if the strength of competition between species is given by Σ_spp_, as might be the case if closely related species are more likely to share common resources, then the relative abundances of species will have covariance matrix (Σ_spp_)^−1^.

Equation 1 is the same as model I in Ives & Helmus (2011), except model I includes variation among species in mean log abundance across sites as fixed effects rather than two random effects, *a*_spp[_*_i_*_]_ and *b*_spp[_*_i_*_]_. This change allows us to align equation 1 with equation 3 (below) that includes variation in the relationship between trait values and log abundance within sites as random effects. In our analyses, treating variation among species in mean log abundance as fixed effects (results not presented) led to almost identical estimates of phylogenetic signal (estimates of 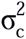and therefore our treatment of *a*_spp[_*_i_*_]_ and *b*_spp[_*_i_*_]_ as random effects does not change the conclusions.

We fit the PLMM with maximum likelihood using function `communityPGLMM` in the `pez`(Pearse *et al*., 2015) package of `R` (R Core Team, 2015). Statistical significance of the variance estimates σ^2^ was determined using a likelihood ratio test. Because the null hypothesis σ^2^ = 0 is on the boundary of the parameter space (σ^2^ cannot be negative), we used the 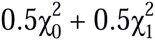 mixture distribution of Self & Liang (1987) for significance tests. The distribution of 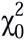 represents a distribution with a point mass at 0, and the *p*-values given by the constrained likelihood ratio test are one-half the values that would be calculated from a standard likelihood ratio test using 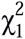. Simulations suggest that *p*-values calculated in this way are more conservative (have higher values) than those from a parametric bootstrap (Appendix Text S1).

Our data set contained many zeros (Fig. 2), raising the question of the validity of applying a linear model to transformed data. Nonetheless, transforming data and applying a linear analysis is robust when assessing the significance of regression parameters (Ives, 2015).

### Can functional traits explain phylogenetic community composition?

To quantify how much of the variation in phylogenetic patterns can be explained by measured functional traits, we estimated PLMMs with and without functional traits, and then compared the strength of phylogenetic signal in the residual variation: if functional traits alone serve to explain phylogenetic community composition, then as functional traits are included, the strength of the phylogenetic signal in the residuals should decrease. We selected functional traits one by one based on the two conditions necessary for them to generate phylogenetic signal in community composition. First, a functional trait must show phylogenetic signal among species, because in the absence of phylogenetic signal among species, a trait could not produce phylogenetic signal in species’ abundances. Second, there must be variation among sites in the relationship between species trait values and abundances; if a trait has phylogenetic signal but there is no variation in relationships between plant functional trait values and abundances among sites, then it will contribute to the overall phylogenetic signal of species abundance and will be captured by *b*_spp[_*_i_*_]_ in equation 1, but it will not affect phylogenetic co-occurrence patterns captured by *c_i_*. Therefore, we only investigate traits that exhibit both strong phylogenetic signal and variation among sites in the apparent advantages the traits give to species.

We tested the phylogenetic signal for each functional trait using model-based methods. Each continuous trait was tested with Pagel’s *λ* (Pagel, 1999) using `phylolm` (Ho & Ané, 2014). For the binary traits, we applied phylogenetic logistic regression (Ives & Garland, 2010) as implemented by `phylolm` (Ho & Ané, 2014). We also tested phylogenetic signal of functional traits via Blomberg's *K* (Blomberg *et al*. 2003) with `picante` (Kembel *et al*. 2010).

We tested variation of relationships between trait values and log abundances with the LMM

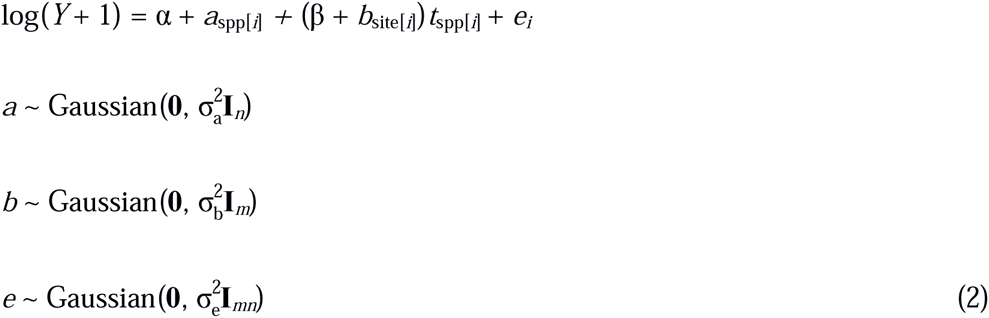

where *t*_spp[_*_i_*_]_ is the focal functional trait value of the species corresponding to observation *i*, and 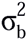 gives the variation among sites in the relationship between species trait values and log abundances. This formulation is closely related to the model used by Pollock *et al*. (2012). If 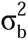, we conclude that different sites select species differently based on the tested trait. We use p < 0.1 here to lower the risk of excluding potential important functional traits.

We quantified the contribution of a trait to the observed phylogenetic pattern in community composition using the model

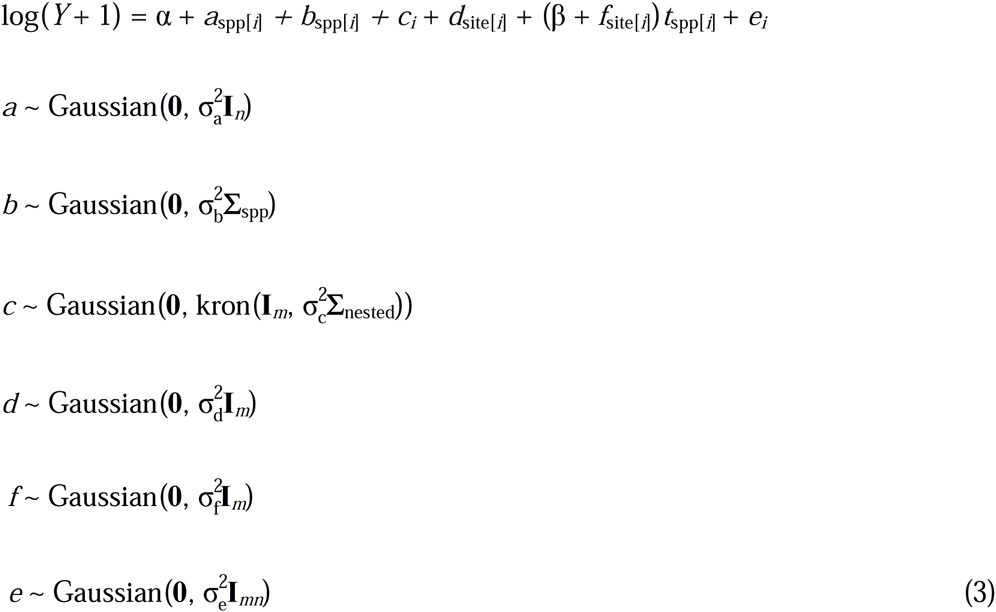

This model is the same as equation 1 used to assess phylogenetic patterns in community composition, except that it includes functional trait values *t*_spp[_*_i_*_]_. The proportion of phylogenetic signal in species composition (estimated by 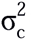) that trait *t*_spp[_*_i_*_]_ can explain is assessed by comparing 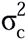 between models with and without this trait as a product with the random effect *f*_site[_*_i_*_]_. Finally, to evaluate the overall contribution of functional traits to the observed phylogenetic patterns, we built a multivariate version of equation 3 which included all traits that have both phylogenetic signal and strong variation among sites.

### Does any environmental variable drive phylogenetic pattern?

If phylogenetic patterns in community composition are observed, yet no functional traits can explain the patterns, how could we identify additional functional traits that might be responsible? Phylogenetically related species usually are assumed to be ecologically similar due to niche conservatism (Wiens *et al*. 2010). Therefore, related species will tend to have similar responses to environmental variables. If these environmental variables are strong enough to drive phylogenetic patterns in community composition, then functional traits that are associated with tolerance or sensitivity to these environmental variables will likely be important in explaining community composition. Thus, we investigated phylogenetic patterns in the responses of species to environmental variables to suggest additional, unmeasured functional traits that might be important to explain phylogenetic patterns in community composition.

We tested for phylogenetic patterns in the responses of species to environmental variables using the PLMM

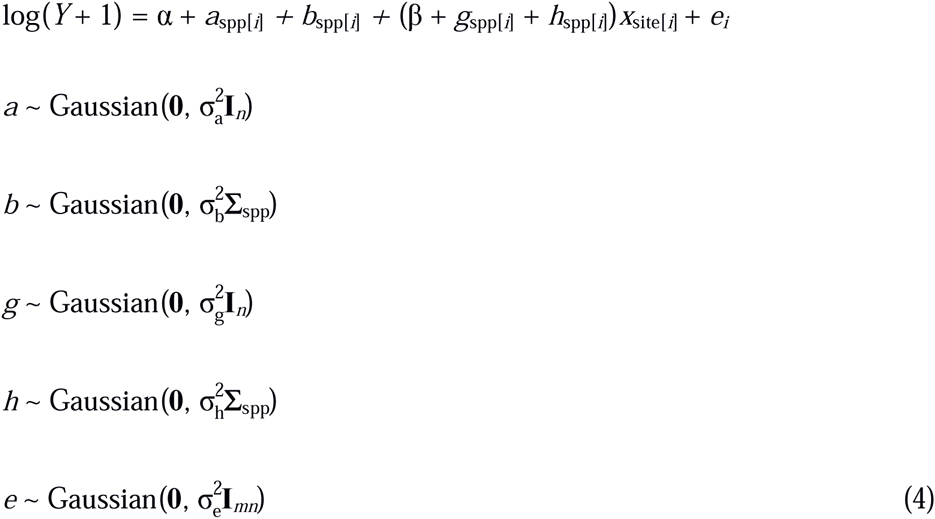

Here, *g*_spp[_*_i_*_]_ and *h*_spp[_*_i_*_]_ represent non-phylogenetic and phylogenetic variation among species in their response to environmental variable *x* (see model II in Ives & Helmus, 2011). The key parameter of interest is 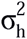, which we tested using a likelihood ratio test. If 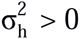, phylogenetically related species respond to environmental variable *x* in similar ways, suggesting the existence of an unmeasured phylogenetically inherited trait that is associated with species tolerances or sensitivities to *x*. Given the large number of environmental variables in our data set, we first applied equation 4 without the term *b*_spp[_*_i_*_]_ and *h*_spp[_*_i_*_]_, and selected environmental variables for which there was variation in responses among species given by *g*_spp[_*_i_*_]_ regardless of whether this variation was phylogenetic. For variables *x* for which 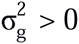 in the reduced version of equation 4, we then applied the full equation 4 and tested whether 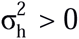.

## Results

### Phylogenetic community composition

Phylogenetically related species co-occurred more often than expected by chance in pine barrens communities in central Wisconsin (Fig. 2). Permutation tests including all 152 species showed that closely related species are likely to have positive covariances in abundance among communities, as judged by either phylogenetic species evenness 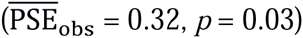 or mean phylogenetic distance 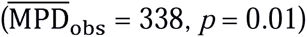. In contrast, when we confine analyses to the 55 focal species that occurring in at least three communities, the permutation tests failed to show statistically significant phylogenetic patterns (abundance data: 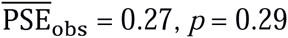; 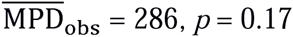; presence/absence data: 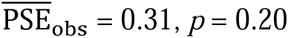; 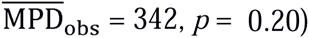). Nevertheless, the PLMM (*p* = 0.008; Table 1) and PGLMM (*p* < 0.001; Appendix Table S1) both reveal statistically significant phylogenetic patterns for the 55 focal species.

**Table 1.**
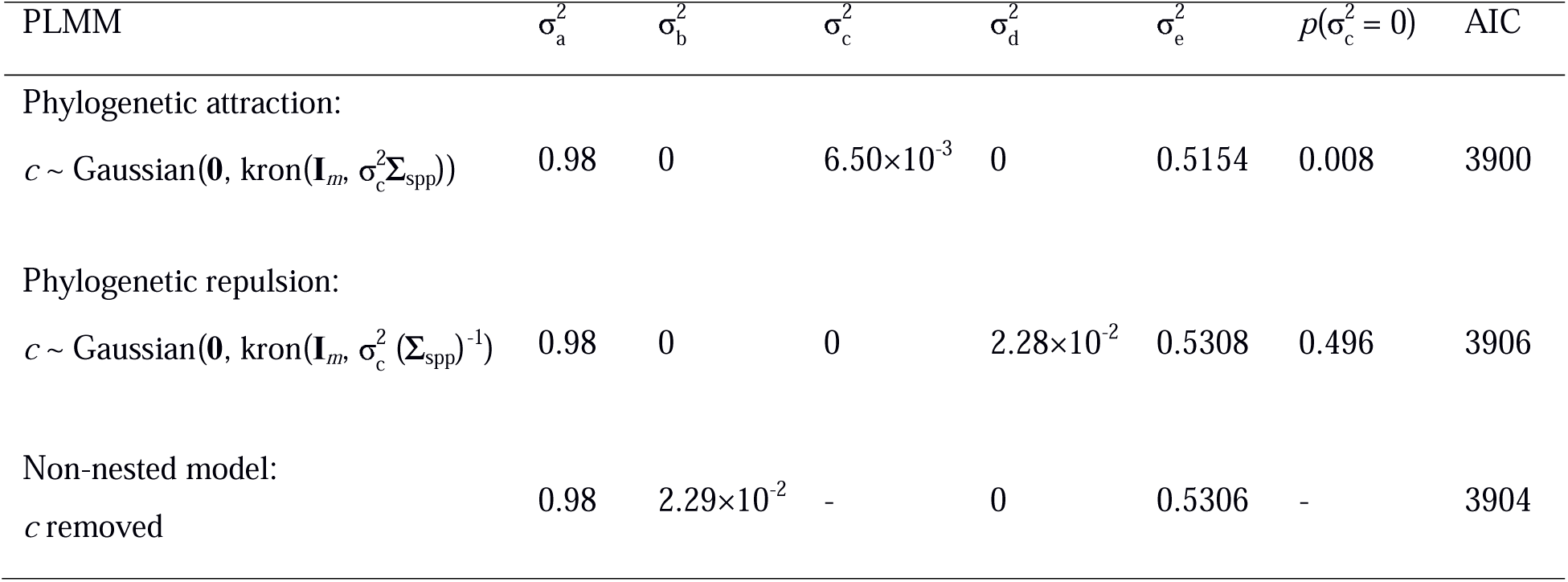
Estimated variance of random effects for the PLMM (equation 1) used to detect phylogenetic patterns in community composition.

### Can functional traits explain phylogenetic community composition?

Most functional traits showed strong phylogenetic signal (Table 2). Five traits – leaf width, leaf thickness, SLA, leaf circularity, and animal dispersal (marginally significant) – also significantly affected plant species’ abundances among sites (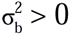, equation 2, Table 2), indicating that different sites selected different species based on these three functional traits. Individually, the five traits reduced the phylogenetic variance in community composition (as measured by reduction in 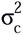 in equation 3 when including these traits) by 18%, 8%, 7%, 2%, and 1%, respectively. Traits that did not pass our two-steps selection individually explained negligible amount of the phylogenetic variance (all <1% and mostly ~0%, data not shown), verifying our initial selection of traits. Including all five traits in the final model reduces the phylogenetic variation 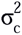 by 57%. Thus, the many functional traits we measured in this study can only reduce the phylogenetic signal in community composition by 57%. Converting the data to presence/absence and using the PGLMM equivalent of equation 3 reduces 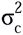 by 89% (Appendix, Table S3). Thus, functional traits explained more of the phylogenetic patterns in the presence/absence of species from communities than in their log abundance, although functional traits still cannot fully explain the phylogenetic pattern in community composition.

**Table 2.**
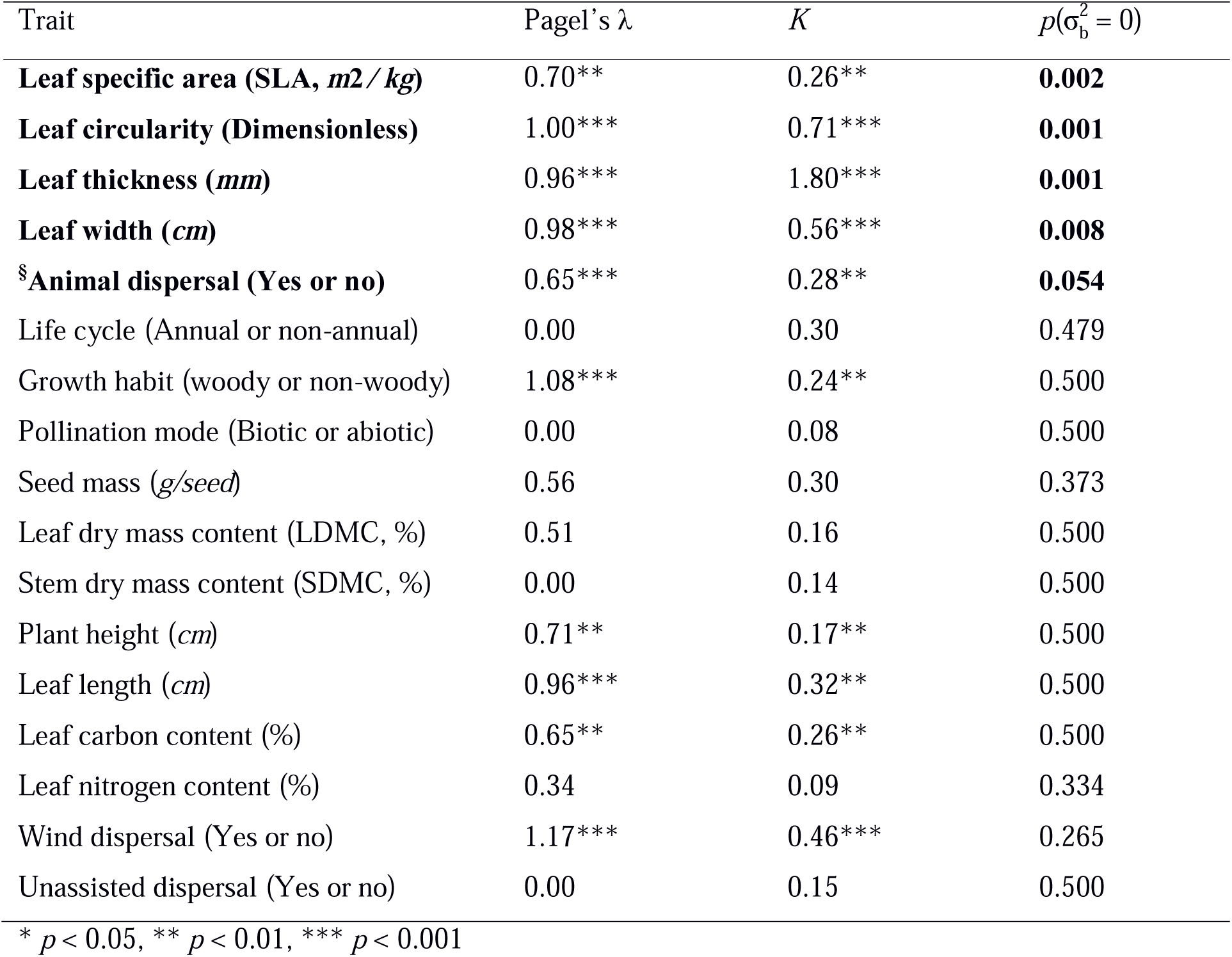
Phylogenetic signal and site variation for each functional trait. P-values for the null hypothesis 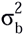 = 0 (equation 2) implying no difference among sites in the effects of trait values on log abundance are given in the column labeled 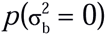. Functional traits with strong phylogenetic signal and 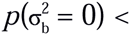 0.1 are considered to be important in explaining phylogenetic patterns.

| Trait | Pagel's $\lambda$ | $K$ | $p(\sigma_b^2 = 0)$ |
| --- | --- | --- | --- |
| **Leaf specific area (SLA, $m^2/kg$)** | 0.70** | 0.26** | **0.002** |
| **Leaf circularity (Dimensionless)** | 1.00*** | 0.71*** | **0.001** |
| **Leaf thickness ($mm$)** | 0.96*** | 1.80*** | **0.001** |
| **Leaf width ($cm$)** | 0.98*** | 0.56*** | **0.008** |
| **§Animal dispersal (Yes or no)** | 0.65*** | 0.28** | **0.054** |
| Life cycle (Annual or non-annual) | 0.00 | 0.30 | 0.479 |
| Growth habit (woody or non-woody) | 1.08*** | 0.24** | 0.500 |
| Pollination mode (Biotic or abiotic) | 0.00 | 0.08 | 0.500 |
| Seed mass ($g/seed$) | 0.56 | 0.30 | 0.373 |
| Leaf dry mass content (LDMC, %) | 0.51 | 0.16 | 0.500 |
| Stem dry mass content (SDMC, %) | 0.00 | 0.14 | 0.500 |
| Plant height ($cm$) | 0.71** | 0.17** | 0.500 |
| Leaf length ($cm$) | 0.96*** | 0.32** | 0.500 |
| Leaf carbon content (%) | 0.65** | 0.26** | 0.500 |
| Leaf nitrogen content (%) | 0.34 | 0.09 | 0.334 |
| Wind dispersal (Yes or no) | 1.17*** | 0.46*** | 0.265 |
| Unassisted dispersal (Yes or no) | 0.00 | 0.15 | 0.500 |

\* $p < 0.05$ , \*\* $p < 0.01$ , \*\*\* $p < 0.001$

**Table 3.** Reduction of the phylogenetic variance in community composition caused by the inclusion of functional traits (equation 3).

| Trait | $\sigma_c^2$ with traits | $\sigma_c^2$ without traits | $100 \times \sigma_{c \text{ (with traits)}}^2 / \sigma_{c \text{ (without traits)}}^2$ |
| --- | --- | --- | --- |
| Leaf width | 0.005302 | 0.006457 | 17.89 |
| Leaf thickness | 0.005921 | 0.006457 | 8.30 |
| SLA | 0.006024 | 0.006457 | 6.71 |
| Leaf circularity | 0.006310 | 0.006457 | 2.28 |
| Animal dispersal | 0.006380 | 0.006456 | 1.18 |
| SLA + circularity + thickness<br>+ Leaf width + Animal<br>dispersal | 0.002804 | 0.006480 | 56.73 |

### Does any environmental variable drive phylogenetic pattern?

There was significant variation among species in their responses to most of the environmental variables we measured, including soil conditions, canopy shade, precipitation, and minimum temperature (Table 4). However, there was no phylogeny signal in the differences among species in their responses to these variables (last column in Table 4). Therefore, no environmental variables we measured can explain the observed phylogenetic pattern in community composition. Using the PGLMM with the presence/absence data, species’ responses to minimum temperature and soil pH, Ca, and Mn concentration all show phylogenetic signal. That is, related species tend to occupy similar sites as measured by these environmental variables (Appendix Table S4). Therefore, functional traits associated with these environmental variables could potentially be responsible for phylogenetic patterns in presence/absence of species among communities.

**Table 4.** Variation in the response of species abundances to environmental variables (equation 4). Although 13/20 environmental variables generated variation in species composition among communities, none of these showed phylogenetic signal in which related species responded more similarly to the environmental variable.

| Environmental variables | *P*-values $\sigma_g^2$ (no<br>phylogenetic signal) | *P*-values for $\sigma_h^2$<br>(phylogenetic signal) |
| --- | --- | --- |
| Minimum temperature | **< 0.001** | 0.500 |
| Precipitation | **< 0.001** | 0.500 |
| Canopy shade | **0.002** | 0.500 |
| Total exchange capacity | **0.002** | 0.500 |
| Organic matter | **0.001** | 0.500 |
| pH | **< 0.001** | 0.500 |
| N | **< 0.001** | 0.500 |
| P | **0.039** | 0.500 |
| Mg | **0.030** | 0.500 |
| K | **0.007** | 0.500 |
| Na | **< 0.001** | 0.500 |
| Mn | **< 0.001** | 0.354 |
| Ca | **< 0.001** | 0.122 |
| Clay | 0.110 | - |
| Silt | 0.070 | - |
| Sand | 0.117 | - |
| Fe | 0.500 | - |
| S | 0.458 | - |
| Zn | 0.500 | - |
| Al | 0.500 | - |

## Discussion

We used our extensive database of functional traits to answer a key question in trait-based and phylogeny-based community ecology: Can information about functional traits explain phylogenetic patterns in community composition? Phylogenetically related plant species are more likely to reach similar abundances in the same pine barren communities of central Wisconsin, yet we could not explain this pattern completely using information about species’ functional traits. When functional traits that themselves showed phylogenetic signal among species were included in the phylogenetic linear mixed model (PLMM) for log abundances of species in communities, that component of the residual variance having phylogenetic covariances decreased by only 57%. The decrease in the phylogenetic component of residual variation was 89% in the analyses of presence/absence data, yet even this leaves residual phylogenetic pattern in the unexplained variation in the presence/absence of species among communities. Thus, even though we measured 15 functional traits, including most of the standard functional traits used to analyze plant community structure, we could not fully explain the phylogenetic patterns in community composition. This suggests that there are either important functional traits that we have not measured, or that there are phylogenetic processes unrelated to functional traits that we have not identified. In either case, these results suggest that including phylogenetic information in addition to functional traits provides further insights into the processes affecting community assembly.

When using the subset of 55 species that occurred in three or more communities, the PLMM (and PGLMM), but not permutation tests, found statistically significant phylogenetic patterns. Ives & Helmus (2011) showed that phylogenetic mixed models have greater statistical power than the metrics like PSE and MPD used with permutation tests. Simulations (Appendix Text S1) show that PLMM analyses tended to have, if anything, incorrectly low Type I error rates, implying that our PLMM results were not the result of false positives. We can thus conclude that closely related species are more likely to co-occur and share similar abundances than expected by chance in these pine barren communities.

Incorporating functional traits reduced the phylogenetic component of residual variation in species composition, what could explain the remaining phylogenetic component? Some unknown historical process might account for this residual phylogenetic variation (Fig. 1B, IV). However, our sites are all located within 100 km with each other, making it unlikely that historical biogeographical processes strongly affect the composition of these communities. It seems more likely that the main source of phylogenetic patterns that were not explained by our measured functional traits is additional unmeasured functional traits. Further analyses of the presence/absence data using PGLMMs suggested that soil conditions (pH, Ca, and Mn levels) and climate (minimum temperature) are potential driving variables for the residual phylogenetic patterns (Appendix Table S3). Traits associated with plant responses to these gradients in environmental conditions could thus account for more of the residual phylogenetic patterns. The functional traits we measured, however, are traits that are unlikely to capture species-specific responses to soil and climatic conditions, and we do not have information on likely traits such as root structure, micorrhizal associations, frost tolerance, etc. We expect such traits might be able to explain more of the phylogenetic pattern in community composition.

We found that functional traits could explain a greater part of the phylogenetic component of the pattern of species presence/absence (89%) than of species abundances (57%). This is unlikely to be a statistical artifact. Because we used only the most common 55 species, detection of species in sites where they occur is likely to be high. In contrast, we expect considerable within-species variation in our estimates of abundance. Because within-species variation will decrease phylogenetic signal (Ives *et al*. 2007), we would expect less residual phylogenetic variation in the abundance data than in the presence/absence data, the opposite of what we found. Therefore, our results suggest that the functional traits we measured have a greater effect on the overall suitability of sites for species than the finer-tuned quality of the sites to support large populations, supporting the argument that including abundance data in phylogenetic community analyses provides more information about community assembly (Freilich & Connolly, 2015).

### Implications

Our results have several implications for community ecology. First, it is clear that studying community composition should incorporate analyses of both phylogenetic structure and functional traits. Phylogenetic and trait information clearly complement each other in allowing sophisticated analyses that can partition the amount of phylogenetic signal in community composition that is associated with functional trait variation (Fig. 1). Our results provide empirical support from community ecology for the argument that phylogenies can provide more information than a set of discretely measured traits (Vane-Wright *et al*. 1991; Cadotte *et al*. 2009). Although functional traits are necessary to accurately infer the processes driving phylogenetic patterns (Kraft *et al*. 2007; Cavender-Bares *et al*. 2009), functional traits alone may often fail to provide a complete picture of community structure.

Second, model-based methods are being increasingly applied in ecology because they are more interpretable, flexible, and powerful than either null models or conventional algorithmic multivariate analyses (Warton *et al*. 2014). With phylogenetic linear mixed models (PLMM), we not only detected phylogenetic patterns in community composition, but also assessed the extent to which these could be explained by functional traits. The ability to combine both phylogenies and functional traits into the same statistical model using PLMMs (and PGLMMs) provides an integrated and quantitative framework for analyzing ecological communities and predicting abundance of one taxon from others.

Finally, we can use phylogenetic analyses to suggest possible unmeasured functional traits that underlie patterns in community composition and that therefore should be measured. If species respond differently to an environmental variable, and if these differences are phylogenetic (i.e., related species respond to the environmental variable in similar ways), then there is likely to be a functional trait or traits that underlie the response of species to this environmental variable. In our study, the phylogenetic patterns in species responses to edaphic conditions like soil chemistry highlighted our lack of data on the specific functional traits related to roots or water/nutrient uptake. While this reveals that our study is incomplete, it also provides a valuable lesson and demonstrates the power of the integrated PLMM approach.

## Acknowledgements

We thank K. Cameron, R. Kriebel, M. Pace, D. Spalink, P. Li, and K. Sytsma for building and providing the phylogeny we used in this study. This project was funded by US-NSF grant DEB-1046355 and DEB-1240804.

## Appendix

In the Appendix we give Tables S1-S4 that correspond to Tables 1–4 in the main text, but using a PGLMM for presence/absence data. The equations used for the PGLMM are the same as equations 1–4, but for binomial data; for example, the PGLMM corresponding to equation 1 is

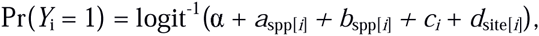

with other terms identical.

**Table S1.** Estimated variance of random effects within the phylogenetic generalized linear mixed model used to detect phylogenetic patterns comparable to equation 1, where phylogenetic attraction and phylogenetic repulsion are estimated by 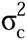.

| PGLMM | $\sigma_a^2$ | $\sigma_b^2$ | $\sigma_c^2$ | $\sigma_d^2$ | $p(\sigma_c^2 = 0)$ |
| --- | --- | --- | --- | --- | --- |
| Phylogenetic attraction: |  |  |  |  |  |
| $c \sim \text{Gaussian}(\mathbf{0}, \text{kron}(\mathbf{I}_m, \sigma_c^2 \mathbf{\Sigma}_{\text{spp}}))$ | 2.84 | 0 | 0.0452 | 0.01 | <0.001 |
| Phylogenetic repulsion: |  |  |  |  |  |
| $c \sim \text{Gaussian}(\mathbf{0}, \text{kron}(\mathbf{I}_m, \sigma_c^2 (\mathbf{\Sigma}_{\text{spp}})^{-1}))$ | 3.10 | 0 | 0.0011 | 0.19 | 0.5 |
| Non-nested model: $c$ removed | 2.83 | 0 | - | 0.18 | - |

**Table S2.**
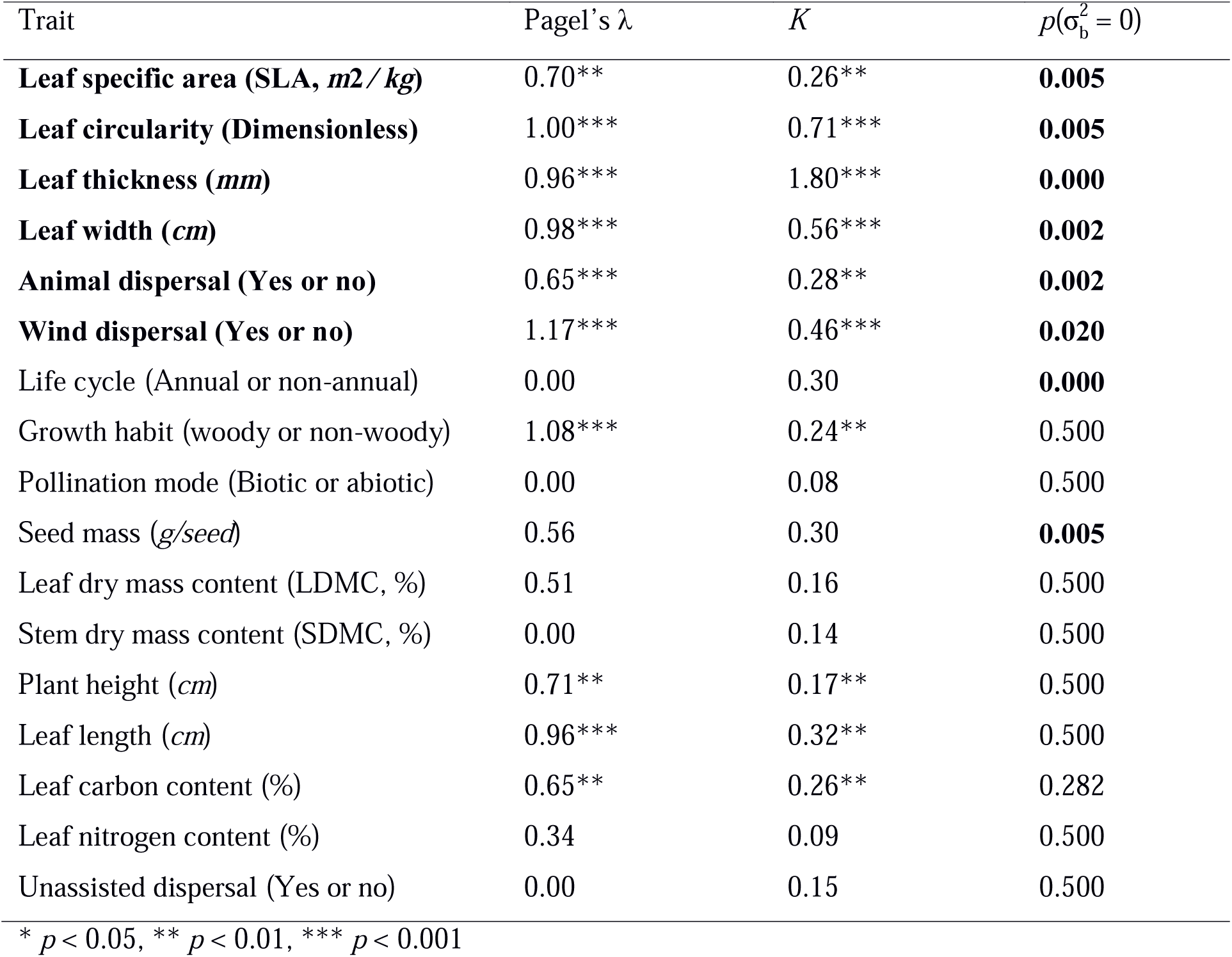
Phylogenetic signal and site variation for each functional trait. P-values for the null hypothesis 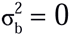 (equation 2) implying no difference among sites in the effects of trait values on presence/absence are given in the column labeled 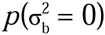. Functional traits with strong phylogenetic signal and 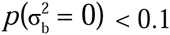 are considered to be important in explaining phylogenetic patterns.

**Table S3.** Proportion of phylogenetic signal of species composition in communities explained by individual functional trait and multiple functional traits. With selected multiple functional traits, about 61% percent of phylogenetic variation was explained, suggesting that phylogenies can provide additional information about community assembly beyond measured functional traits. See equation 3 in the Methods section for details about models.

| Trait | $\sigma_c^2$ with trait | $\sigma_c^2$ without trait | $100 \times \sigma_c^2(\text{with trait}) / \sigma_c^2(\text{without trait})$ |
| --- | --- | --- | --- |
| Leaf width | 0.018105 | 0.041847 | 56.74 |
| Leaf thickness | 0.030925 | 0.041847 | 26.10 |
| Leaf circularity | 0.035442 | 0.041854 | 15.32 |
| SLA | 0.036811 | 0.041828 | 12.00 |
| Wind dispersal | 0.039946 | 0.041844 | 4.54 |
| Animal dispersal | 0.041534 | 0.041862 | 0.78 |
| Leaf width + Leaf thickness +<br>Leaf circularity + SLA + Wind<br>dispersal + Animal dispersal | 0.004616 | 0.041834 | 88.97 |

**Table S4.** There are strong variations in species’ relationships between their presence/absence and most environmental variables (*p* value of each environmental variable was presented in the *P*-values for variation column). Four of these variations show phylogenetic signal. For environmental variable that has no strong variation in species’ responses, no further test for phylogenetic signal of variation was conducted (thus “-” in the third column). P-values that are less than 0.05 are in bold.

| Environmental variables | P-values for variation | P-values for phylogenetic signal of variation |
| --- | --- | --- |
| **Minimum temperature** | **<0.001** | **0.002** |
| Precipitation | **<0.001** | 0.500 |
| Canopy shade | **0.001** | 0.500 |
| Total exchange capacity | 0.149 | - |
| Organic matter | 0.161 | - |
| **pH** | **0.005** | **<0.001** |
| N | 0.052 | - |
| P | 0.343 | - |
| Mg | 0.500 | - |
| K | 0.206 | - |
| Na | **0.004** | 0.500 |
| **Mn** | **<0.001** | **<0.001** |
| **Ca** | **0.012** | **<0.001** |
| Clay | 0.431 | - |
| Silt | 0.494 | - |
| Sand | 0.500 | - |
| Fe | 0.379 | - |
| S | 0.500 | - |
| Zn | 0.500 | - |
| Al | 0.500 | - |

**Text S1.**
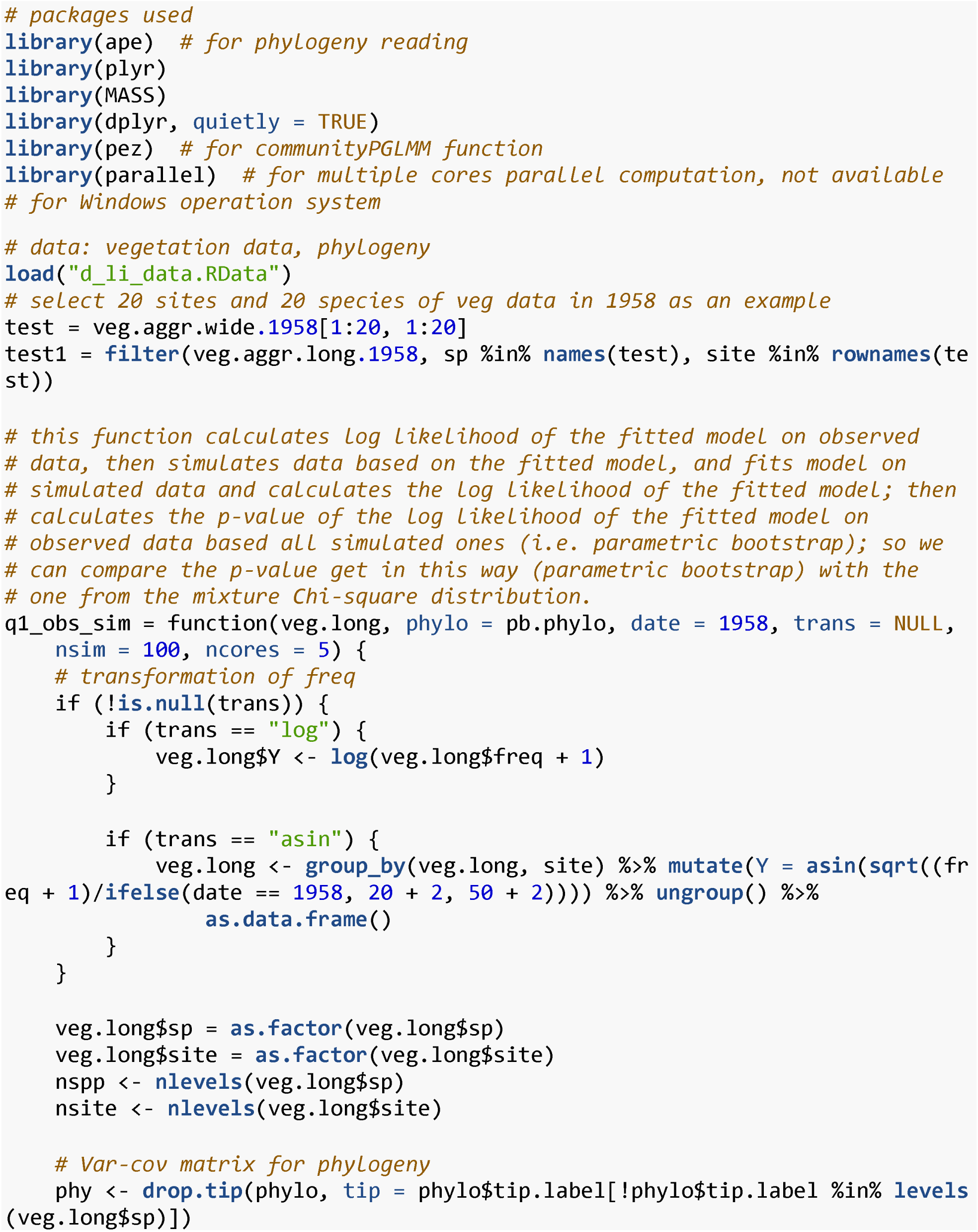
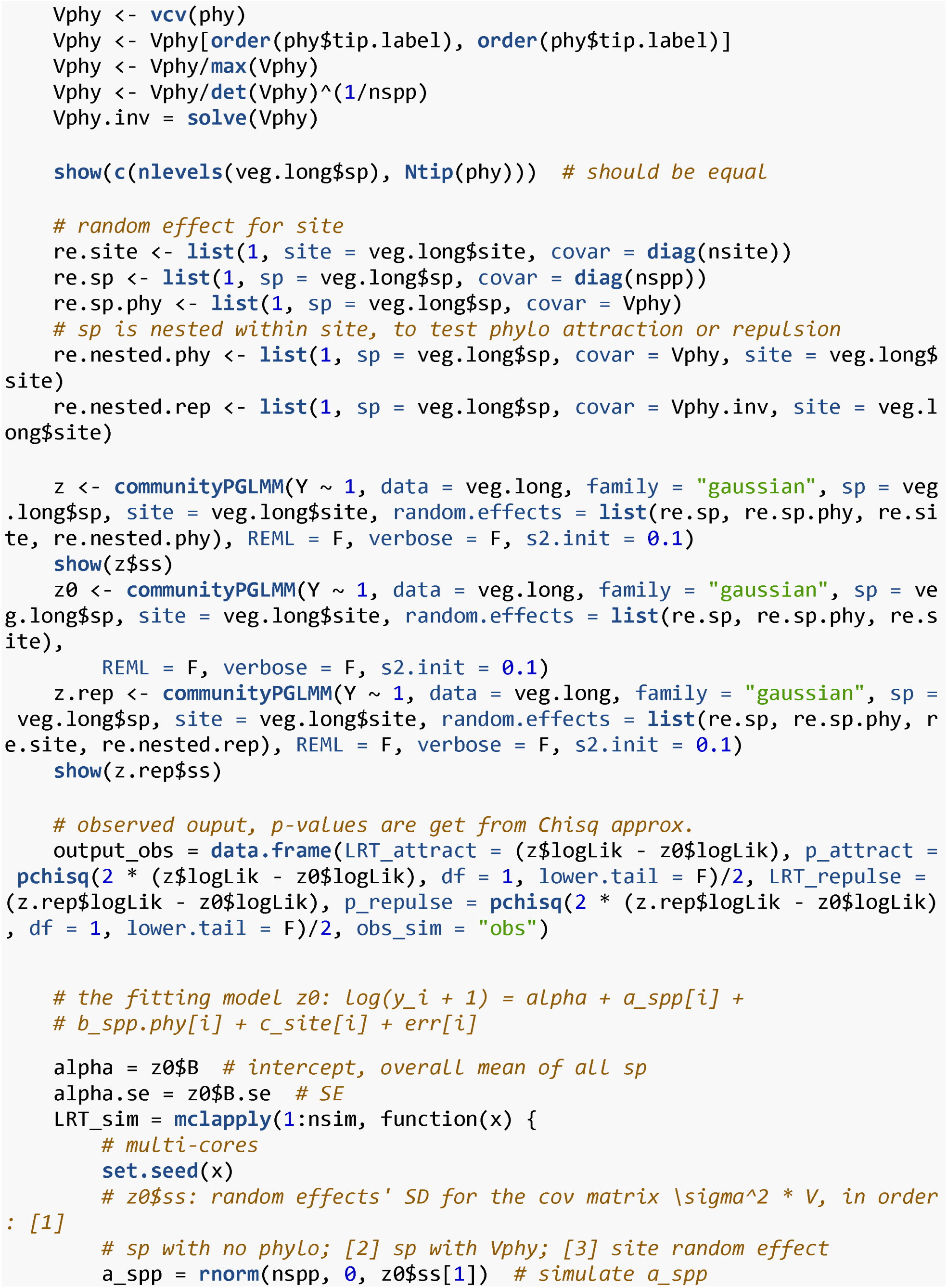
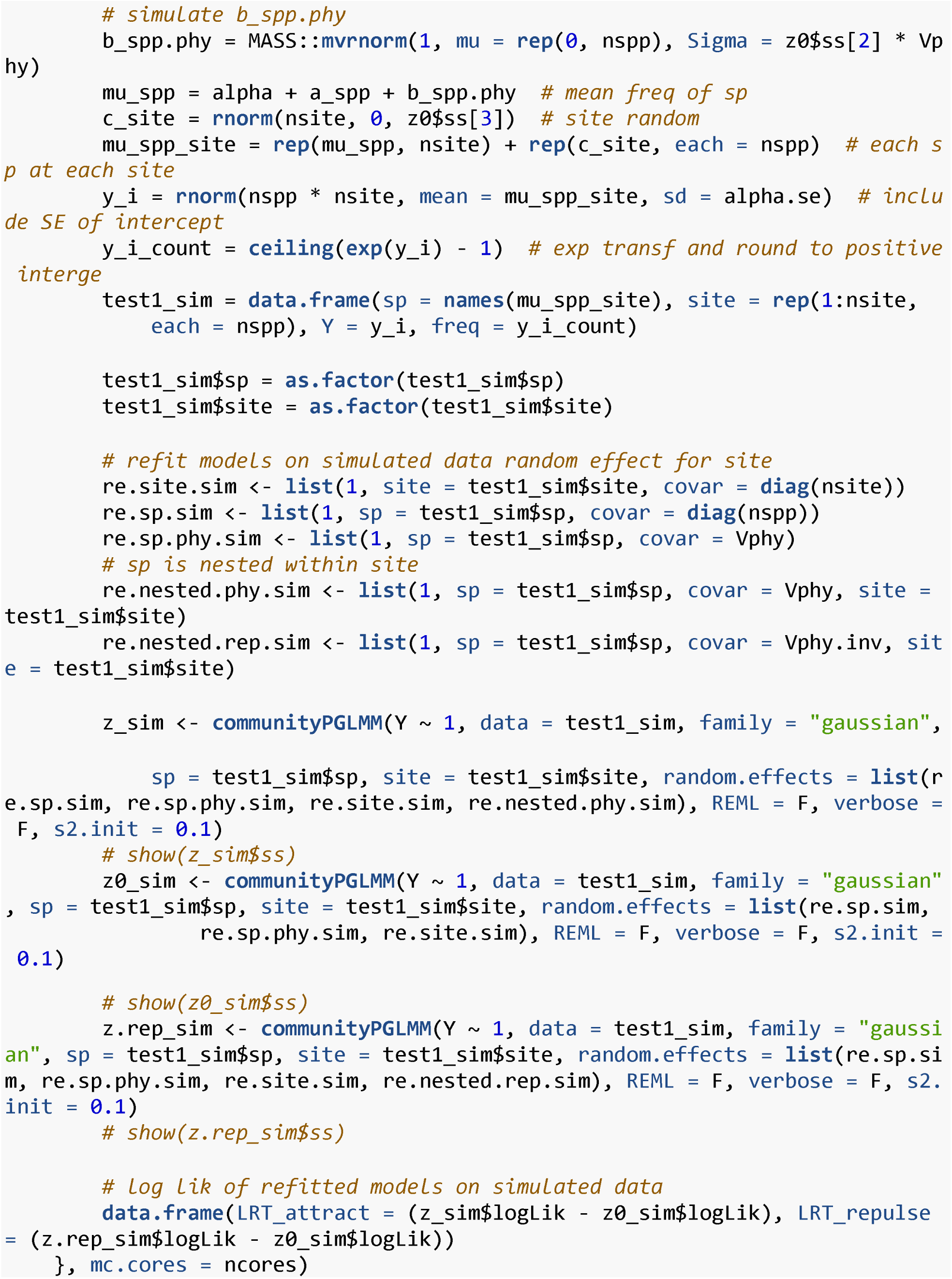
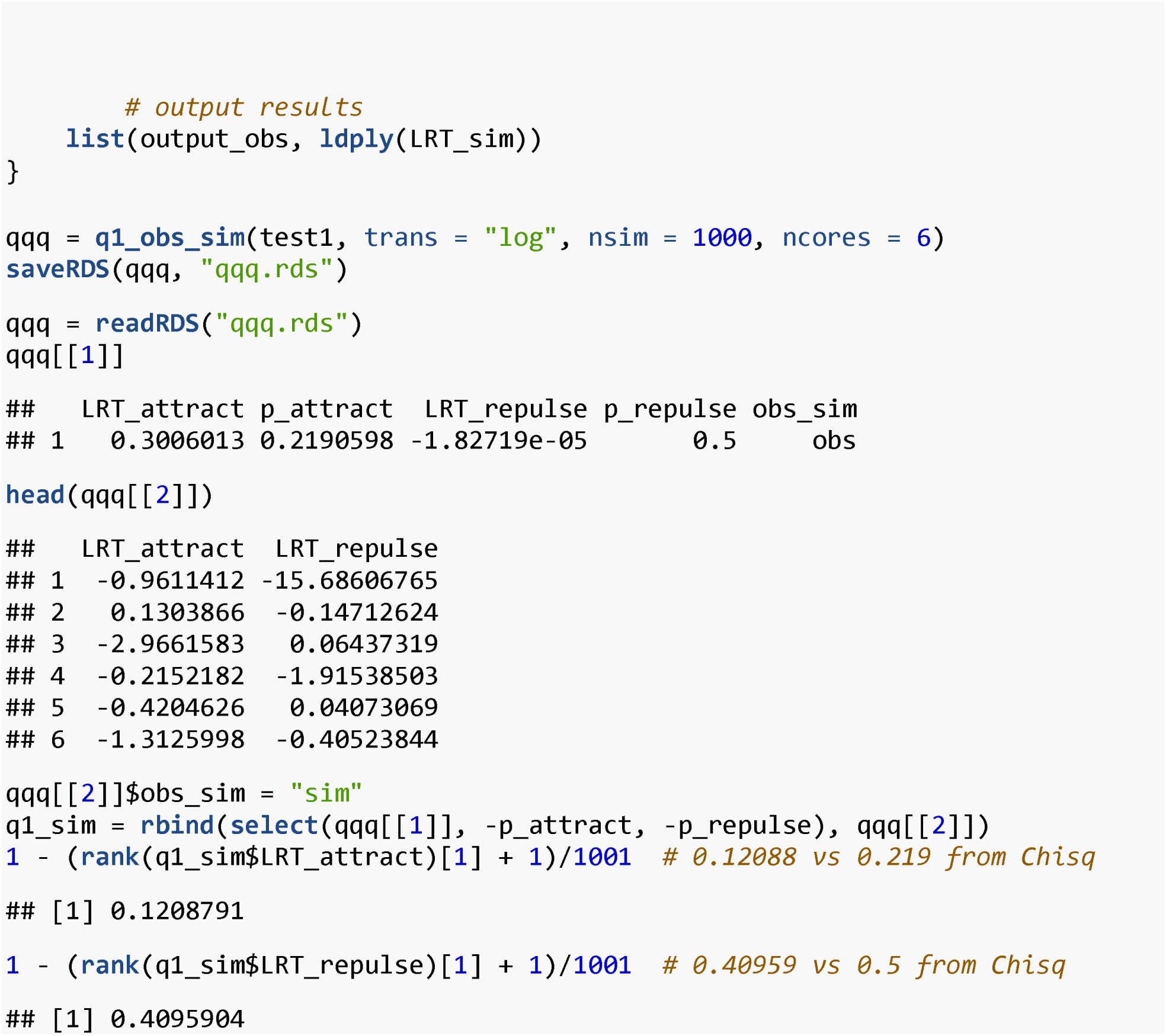
Code to compare *p*-values of null hypothesis σ^2^ = 0 calculated from the 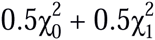 mixture distribution and parametric bootstrap. The *p*-values based on the mixture Chi-square distribution are conservative (i.e. higher than those from parametric bootstrap).

## Footnotes

1 Cameron, K., R. Kriebel, M. Pace, D. Spalink, P. Li, and K. Sytsma. *In prep*. A complete molecular community phylogeny for the flora of Wisconsin based on the universal plant DNA barcode.

